# Asymmetric assembly of congested contiguous stereocenters via photobiocatalytic three-component radical coupling

**DOI:** 10.64898/2026.09.25.754468

**Authors:** Qingdong Hu, Cang-Xin Zheng, Binh Khanh Mai, Peng Liu, Yang Yang

**Affiliations:** Department of Chemistry and Biochemistry, University of California, Santa Barbara, California 93106, USA; Department of Chemistry, University of Pittsburgh, Pittsburgh, Pennsylvania 15260, USA; Department of Bioengineering, University of California, Santa Barbara, California 93106, USA; Biomolecular Science and Engineering (BMSE) Program, University of California, Santa Barbara, California 93106, USA; Howard Hughes Medical Institute, University of California, Santa Barbara, California 93106, USA

## Abstract

Despite rapid advances in new-to-nature biocatalysis, trimolecular enzymatic reactions that generate previously inaccessible molecular entities, particularly those bearing well-defined, congested stereochemical dyads, remain exceedingly rare. Here, we report a novel diastereo- and enantioselective photobiocatalytic three-component radical coupling unknown in both organic chemistry and enzymology, enabled by cooperative catalysis employing an evolved pyridoxal biocatalyst and a transition-metal photosensitizer. Directed evolution using high-throughput photobiocatalysis furnished two threonine aldolase variants, enabling the stereoselective assembly of contiguous tri- and tetrasubstituted stereocenters as well as vicinal tetrasubstituted stereocenters, which were long regarded as a challenging goal for stereoselective radical chemistry and asymmetric catalysis. Furthermore, a variety of radical precursors, including α-iodoesters, α-iodoamides, α-iodoketones, and α-iodonitriles, could be transformed into structurally diverse, densely substituted non-canonical amino acid derivatives via C–H functionalization with excellent stereocontrol. Leveraging the broad substrate compatibility of engineered pyridoxal biocatalysts, combinatorial biocatalysis was achieved by simultaneously varying all three coupling partners, delivering products with a 98% success rate. Collectively, this diversity-generating multicomponent photobiocatalytic radical coupling afforded a powerful strategy for addressing long-standing challenges in asymmetric catalysis while enabling access to valuable stereochemically complex small molecules with applications in medicinal chemistry.

---

Although the past decade has witnessed considerable progress in unnatural photobiocatalysis^1–23^, the rational design and development of enzymatic processes which are not only new to nature but also unknown to synthetic organic chemistry remain a central challenge^10^. Reprogramming of organic cofactor-dependent^1–15^ and independent^16–18^ enzymes as well as metalloenzymes^19–23^ has delivered elegant solutions for catalytic stereocontrol over radical intermediates; however, the vast majority of previously reported biocatalytic reactivity mirrors transformations previously established in synthetic chemistry using small-molecule catalysts^1–5,13–20, 22–23^. Consequently, the identification of general enzymatic strategies for accomplishing chemically novel and synthetically valuable catalytic reactions has emerged as a key frontier in the rapidly evolving field of enzyme reprogramming and directed evolution^24, 6–12, 21^.

While unimolecular and bimolecular photobiocatalytic reactions have become common after a decade of intensive research, trimolecular photobiocatalytic coupling processes have only begun to emerge^25,10,15^. In native enzymology, few single-domain natural enzymes are capable of simultaneously engaging three distinct substrates, rendering naturally occurring trimolecular enzymatic reactions exceedingly rare. Instead, nature uses multi-domain megaenzyme complexes such as polyketide synthases (PKSs)^26^ and nonribosomal peptide synthetases (NRPSs)^27^ or multienzyme cascades^28^ to orchestrate complex multicomponent biotransformations. By harnessing synthetic photochemistry to generate reactive intermediates via intermolecular processes, photobiocatalysis circumvents the stringent requirement for trimolecular substrate binding, thereby enabling enzymatic multicomponent coupling reactions that extend beyond the reactivity landscape of natural enzymes. Guided by this concept, we recently demonstrated stereoselective three-component reactions using pyridoxal β-decarboxylases through a decarboxylative β-C–C coupling mechanism^10^. Building on the mechanistic and structural diversity of pyridoxal 5’-phosphate (PLP) dependent enzymes, we sought to generalize photobiocatalytic three-component coupling to other PLP activation mechanisms for tackling long-standing challenges in stereoselective radical chemistry and asymmetric catalysis, particularly the construction of congested stereogenic centers.

In this context, we recently questioned whether α-functionalizing pyridoxal enzymes could be leveraged to achieve a novel class of photobiocatalytic three-component C–C coupling between organic halides (**1**), vinylarenes (**2**) and amino acids (**3**) via an α-C–H functionalization logic (Fig. 1a). We further recognized that such an enzymatic transformation could provide diastereo- and enantioselective access to sterically congested contiguous stereogenic centers^29–31^, including vicinal tri- and tetrasubstituted stereocenters as well as two vicinal tetrasubstituted stereocenters^30,31^, a process that continues to present significant hurdles for stereoselective catalysis, particularly those involving a free radical mechanism^29^ (Fig. 1b).

**Fig. 1.**
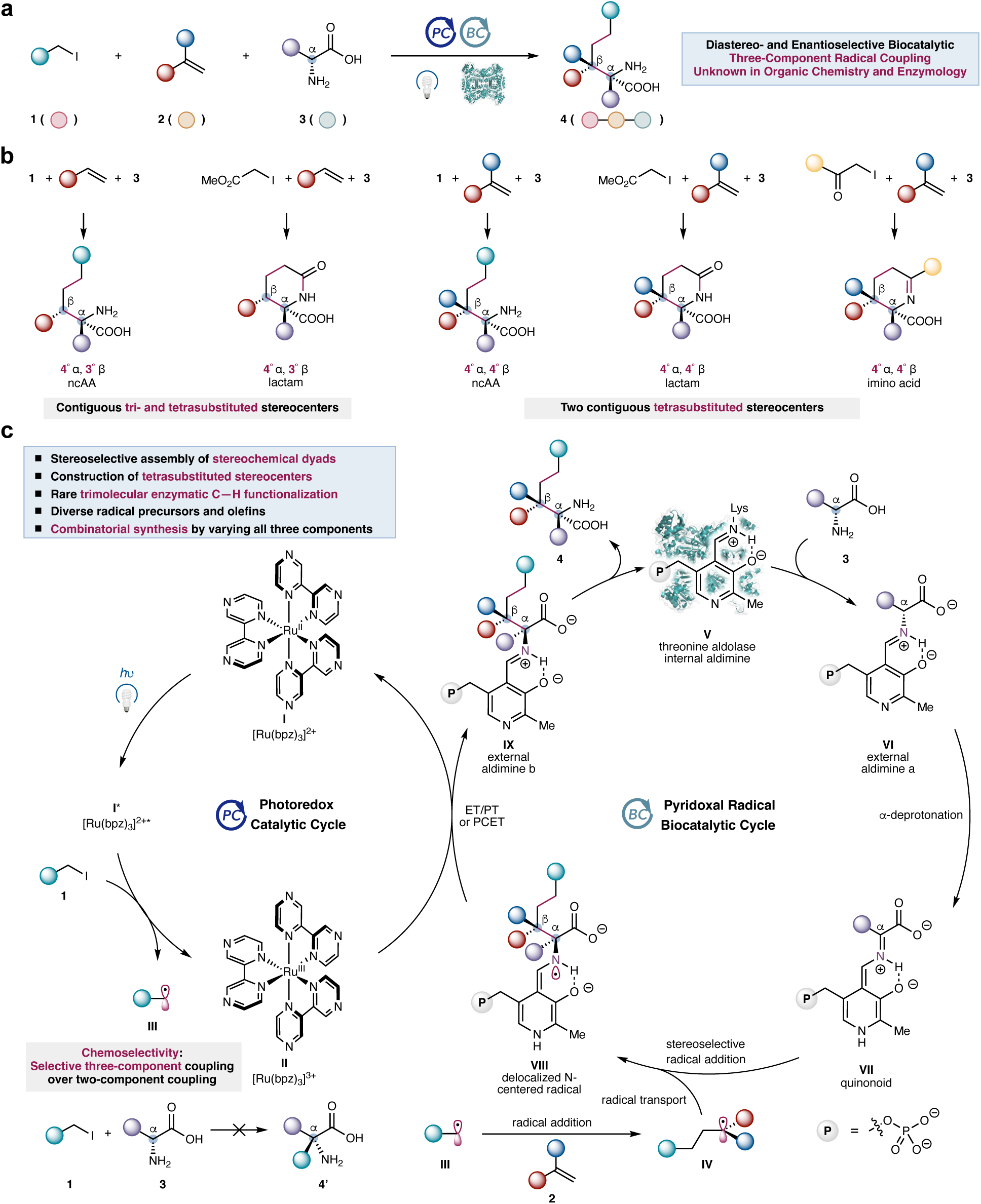
Photobiocatalytic three-component coupling for the diastereo- and enantioselective construction of challenging stereochemical dyads. **a**, Threonine aldolase-catalyzed stereoselective three-component coupling of α-halo compounds, vinylarenes and abundant amino acid substrates: overall transformation. b, Diastereo- and enantioselective assembly of contiguous tri- and tetrasubstituted stereocenters as well as contiguous tetrasubstituted stereocenters via photobiocatalytic three-component coupling. c, Proposed dual catalytic cycles for the photobiocatalytic three-component coupling. ncAA: non-canonical amino acid.

Our proposed dual photobiocatalytic cycle is detailed in Fig. 1c. In the photoredox cycle, excitation of a suitable photosensitizer (**I**), such as Ru(bpz)_3_Cl_2_, would allow photoinduced electron transfer to α-halo organic substrate (**1**), generating an electrophilic radical species (**III**). Before entering the enzyme active site, this nascent radical would be intercepted by a vinylarene substrate (**2**), furnishing a nucleophilic benzylic radical (**IV**) poised for enzymatic capture. Concurrently, in the biocatalytic cycle, transimination of the PLP enzyme internal aldimine (**V**) with the amino acid donor (**3**) would afford an external aldimine **VI**. Subsequent α-deprotonation of **VI** mediated by an active-site basic residue would generate a persistent enzymatic quinonoid intermediate (**VII**). The photocatalytically formed benzylic radical (**IV**) would then diffuse into the active site and add to the Cα position of the quinonoid intermediate in an enzyme-controlled diastereo- and enantioselective fashion. This C–C bond forming radical addition would give rise to a delocalized N-centered radical (**VIII**), which would be subsequently oxidized by the oxidized form of the ground-state photocatalyst (**II**) to form a new external aldimine intermediate (**IX**). Final transimination with the conserved lysine residue of the PLP enzyme would release the stereochemically well defined three-component coupling product (**4**) and regenerate the biocatalyst, thereby completing the catalytic cycle.

To realize this proposed three-component coupling, we would need to effectively suppress the competing two-component coupling between the α-halo substrate **1** and amino acid substrate **3** (**1** + **3** → **4’**, Fig. 1c). We hypothesized that this chemoselectivity could be accomplished by accelerating the interception of the transient radical intermediate **III** with olefin **2** while simultaneously promoting addition of the nascent radical **IV** to the enzymatic quinonoid **VII** through the judicious selection and engineering of α-functionalizing PLP enzymes. We further envisioned that a wide range of α-halo compounds (**1**), including α-haloesters, α-haloamides, α-haloketones and α-halonitriles, could be rendered compatible with this photobiocatalytic three-component coupling upon directed evolution. Diverse vinylarenes (**2**), including α-substituted derivatives, could also be engaged with this dual catalytic activation manifold, allowing the generation and stereocontrolled transformation of both secondary and tertiary benzylic radicals. Moreover, both glycine and α-branched amino acid donors could be transformed via a C–H functionalization logic. Collectively, these features would establish this photobiocatalytic three-component coupling as a general platform for the stereocontrolled construction of sterically congested contiguous stereogenic centers, including vicinal tetrasubstituted carbons and vicinal tri- and tetrasubstituted carbons. Herein, we describe the successful implementation of this proposed photobiocatalytic three-component coupling.

## Results and discussion

### Discovery and directed evolution of three-component coupling biocatalysts for the construction of contiguous tri- and tetrasubstituted stereocenters

At the outset of this study, we sought to identify a suitable PLP-dependent biocatalyst capable of facilitating the proposed photobiocatalytic three-component coupling of an α-iodoacetate (**1a**), styrene (**2a**), and alanine (**3a**) to access non-canonical amino acids (ncAAs) bearing contiguous tri- and tetrasubstituted stereocenters (Fig. 2). Using Ru(bpy)_3_Cl_2_ as the photoredox catalyst, we surveyed a panel of PLP enzymes catalyzing the formation of C–C bonds at the α-position of amino acids (Fig. 2a). Although no coupling product **4a** was detected with the biosynthetic PLP enzymes ObiH^32,33,34,35^ and LolT^36^ and PLP-dependent serine hydroxymethyltransferases from *Eschericia coli* (*Ec*SHMT)^37^, low but measurable three-component coupling activity was observed within the threonine aldolase family. When 1 mol% threonine aldolase homologs were employed, the photobiocatalytic reaction furnished the desired three-component coupling product **4a** along with lactam **5a**, arising from spontaneous lactamization of **4a**. To enable accurate determination of yield and diastereoselectivity, the biocatalytic reaction mixture was incubated under mildly basic conditions (pH = 10) at 60 °C, effecting clean conversion of amino ester **4a** into lactam **5a**. Under these conditions, in the presence of 2 mol% Ru(bpy)_3_Cl_2_, *Escherichia coli* threonine aldolase (*Ec*TA)^38^ and *Aeromonas jandaei* threonine aldolase (*Aj*TA)^39,40^ afforded **5a** in 6% yield (87:13 diastereomeric ratio (d.r.), >99:1 enantiomeric ratio (e.r.)) and 4% yield (85:15 d.r., >99:1 e.r.), respectively. As no alternative chemical synthesis was available to access racemic **5a** and related stereochemical dyads, enantiomeric ratios were as determined by derivatization of the carboxylic acids with chiral amine agent and subsequent HPLC analysis (see Figure S2 for details).

**Fig. 2.**
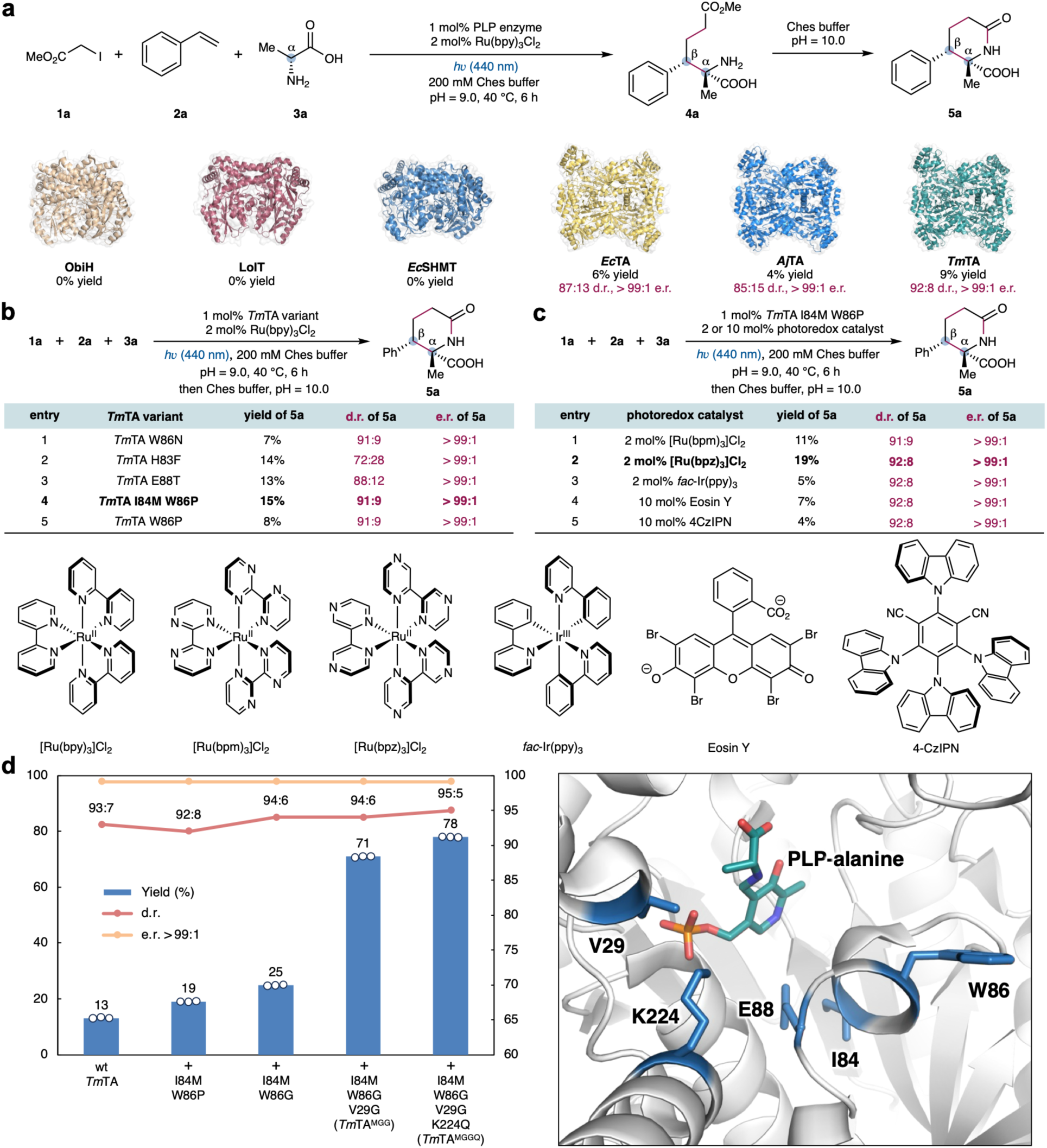
Discovery, optimization and directed evolution of three-component coupling biocatalysts for the construction of contiguous tri- and tetrasubstituted stereocenters. **a**, Evaluation of α-functionalizing pyridoxal enzymes. Reaction conditions for photobiocatalysis: **1a** (2.0 equiv, 8.0 mM), **2a** (1.0 equiv, 4.0 mM), **3a** (5.0 equiv, 20.0 mM), 2 mol% Ru(bpy)_3_Cl_2_ (80 μM), 1 mol% PLP enzyme (40 μM), 200 mM Ches buffer (pH = 9.0), *h*ν (440 nm Kessil LED, 40 W, 100% intensity), 40 °C, 6 h. Lactamization conditions: 400 µL Ches buffer (200 mM, pH = 10.0) was added, 60 °C, 6 h. **b**, Evaluation of engineered *Tm*TA variants from our in-house enzyme collection. **c**, Selected results on the evaluation of photoredox catalysts. **d**, Directed evolution of *Tm*TA I84M W86P leading to the final variant *Tm*TA I84M W86G V29G K224Q (*Tm*TA^MGGQ^). The active-site illustration was made from PDB ID 1LW5^41^.

Among the homologs examined, *Thermotoga maritima* threonine aldolase (*Tm*TA)^41^ exhibited the highest initial activity, affording **5a** in 9% yield with 92:8 d.r. and >99:1 e.r. (Fig. 2a). With this initial result, we next evaluated a focused library of engineered *Tm*TA variants in our enzyme library. Although *Tm*TA W86N previously engineered for oxidative coupling with organoboron reagents^8^ activity compared relative to the wild-type enzyme, *Tm*TA H83F^8^ showed enhanced activity but modest diastereoselectivity (14% yield, 72:28 d.r. and >99:1 e.r., Fig. 2b, entry 2). Similarly, *Tm*TA E88T previously engineered in our laboratory for α-alkylation of amino acids using Katritzky salts^7^ furnished improved activity with slightly diminished diastereocontrol (13% yield, 88:12 d.r. and >99:1 e.r., entry 3). Notably, the double mutant *Tm*TA I84M W86P displayed the highest initial activity and diastereoselectivity (15% yield, 91:9 d.r., >99:1 e.r., entry 4), whereas the single mutant *Tm*TA W86P exhibited activity comparable to that of wt *Tm*TA. These results highlighted the key role of the I84M mutation. *Tm*TA I84M W86P was therefore selected as the template for further optimization. Importantly, two-component coupling product **4a’** derived from **1a** and **2a** was not observed under these reaction conditions, indicating the excellent chemoselectivity favoring the desired three-component coupling.

Using *Tm*TA I84M W86P, we next examined a range of transition-metal and organic photoredox catalysts. Among all the photocatalysts evaluated, Ru-based photosensitizers consistently provided superior results (see Table S3 for details). Ru(bpy)_3_Cl_2_ furnished **5a** in 15% yield with 91:9 d.r. and >99:1 e.r. (Fig. 2b, entry 4). Screening of related Ru complexes revealed that Ru(bpm)_3_Cl_2_ was less effective (11% yield, 91:9 d.r. and >99:1 e.r., Fig. 2c, entry 1), whereas Ru(bpz)_3_Cl_2_ led to improved efficiency, affording **5a** in 19% yield, 92:8 d.r. and >99:1 e.r. (Fig. 2c, entry 2). Other transition-metal photocatalysts, including *fac*-Ir(ppy)_3_, provided lower yields of **5a** (see Table S3 for details). Organic photocatalysts, including Eosin Y (Fig. 2c, entry 4) and rhodamine-based dyes as well as 4CzIPN (Fig. 2c, entry 5), were markedly less effective for this transformation (see Table S3 for details).

With these results in hand, we set out to evolve *Tm*TA I84M W86P to improve both its activity and diastereoselectivity for the photobiocatalytic three-component coupling (Fig. 2d). Guided by the crystal structure of *Tm*TA (PDB ID: 1LW5^41^), we applied site-saturation mutagenesis (SSM) to generate enzyme variant libraries by diversifying active-site residues proximal to the PLP covalent intermediate. For each SSM library, 88 clones were selected for screening. To enable enzyme engineering under photobiocatalytic conditions, we established an in-house high-throughput screening platform. *Tm*TA variants were overexpressed in *Escherichia coli* in 24-well plates, lysed by sonication using a 24-tip horn, and the resulting clarified cell-free lysates were subjected to photobiocatalytic screening. Reactions were performed using a custom-built parallel photoreactor equipped with Kessil LED illumination and analyzed by ultraperformance liquid chromatography–mass spectrometry (UPLC–MS).

We first targeted active-site residue W86 located within the 83–88 α-helix proximal to the PLP cofactor for SSM. From this initial round of engineering, *Tm*TA I84M W86G emerged as an improved variant, showing slightly enhanced activity and diastereoselectivity, providing **5a** in 25% yield, 94:6 d.r., and >99:1 e.r. (Fig. 2d). In contrast to our previous studies on *Tm*TA engineering for radical C–C coupling^7,8^, further saturation of residues within this α-helix did not lead to additional improvements. We thus shifted our focus to N-terminal residues positioned near the Cα of the PLP covalent intermediate. Subsequent SSM and screening revealed V29G as a highly beneficial substitution, resulting in an approximately three-fold increase in catalytic activity while fully preserving diastereoselectivity (71% yield, 94:6 d.r. and >99:1 e.r.). Notably, V29G represented a previously unrecognized yet critical mutation for enabling novel C–C bond forming reactivity in *Tm*TA. A final round of directed evolution targeting residue K224 revealed K224Q as an additional beneficial mutation, providing the final quadruple mutant *Tm*TA I84M W86G V29G K224Q (*Tm*TA^MGGQ^). Under optimized conditions using 1 mol% *Tm*TA^MGGQ^ and 2 mol% Ru(bpz)_3_Cl_2_, three-component coupling product **5a** formed in 78% yield, 95:5 d.r. and >99:1 e.r., corresponding to a six-fold enhancement in activity relative to wild-type *Tm*TA, allowing the diastereo- and enantioselective construction of stereochemical dyads featuring contiguous tri- and tetrasubstituted stereocenters.

Further investigation revealed that _D_-alanine (_D_-**3a**) underwent more efficient conversion than _L_-alanine (_L_-**3a**, see Table S5, entries 1 and 7). Racemic _DL_-alanine (_DL_-**3a**) could also be employed as the amino acid donor with comparable efficiency (Table S5, entry 8). The use of α-bromoacetate in lieu of **1a** afforded only 10% yield of **5a** (Table S5, entry 15), indicating the importance of the choice of α-haloester substrate. Control experiments conducted in the absence of *Tm*TA^MGGQ^ or visible-light irradiation produced no detectable **5a**, thereby confirming the dual photobiocatalytic nature of the transformation (see Table S5 for full details).

### Substrate scope of photobiocatalytic construction of contiguous tri- and tetrasubstituted stereocenters

Having established the optimal reaction conditions, we next investigated the substrate scope of this photobiocatalytic three-component coupling for the stereoselective construction of vicinal tri-and tetrasubstituted stereocenters (Fig. 3). It was found that vinylarenes bearing diverse sensitive *para*-substituents, including a pinacol boronic ester (**5b**), an acetamide bearing an NH moiety (**5c**) and a hydroxy group (**5d**), were readily accommodated, providing the corresponding three-component coupling products with excellent diastereo- and enantioselectivity. Notably, a large *para*-phenyl group (**5e**) was also tolerated, showcasing the broad substrate compatibility of the engineered threonine aldolase biocatalysts. Halogen substituents such as a fluorine (**5f**), a chlorine (**5g**), and a bromine (**5h**) group were also compatible. Additionally, styrenes bearing a *meta*-methoxy (**5i**) and an *ortho*-methyl (**5j**) group underwent efficient coupling with excellent diastereo- and enantioselectivity. Heterocyclic substrates including a 1,2,4-triazole (**5k**) and a thiophene (**5l**) were also successfully transformed into the corresponding stereochemical dyads. Furthermore, without additional enzyme engineering, *Tm*TA^MGGQ^ readily converted α-iodoacetonitrile (**1e**), delivering the corresponding acyclic non-canonical amino acid with adjacent tri- and tetrasubstituted stereocenters (**4m**) in 71% yield, 87:13 d.r. and >99:1 e.r.. Moreover, another *Tm*TA variant *Tm*TA E88V I84A F85R (*Tm*TA^VAR^) was found to efficiently convert glycine (**5n**), allowing the stereoselective construction of two adjacent trisubstituted stereocenters. Importantly, a preparative-scale synthesis of **5a** was achieved on a 147 mg scale with a reduced enzyme loading of 0.5 mol% without compromising the stereoselectivity, demonstrating the practicality of the photobiocatalytic three-component coupling. The relative and absolute stereochemistry of **5f** was unambiguously established by single-crystal X-ray diffraction analysis (CCDC accession number 2500477, see the SI for details).

**Fig. 3.**
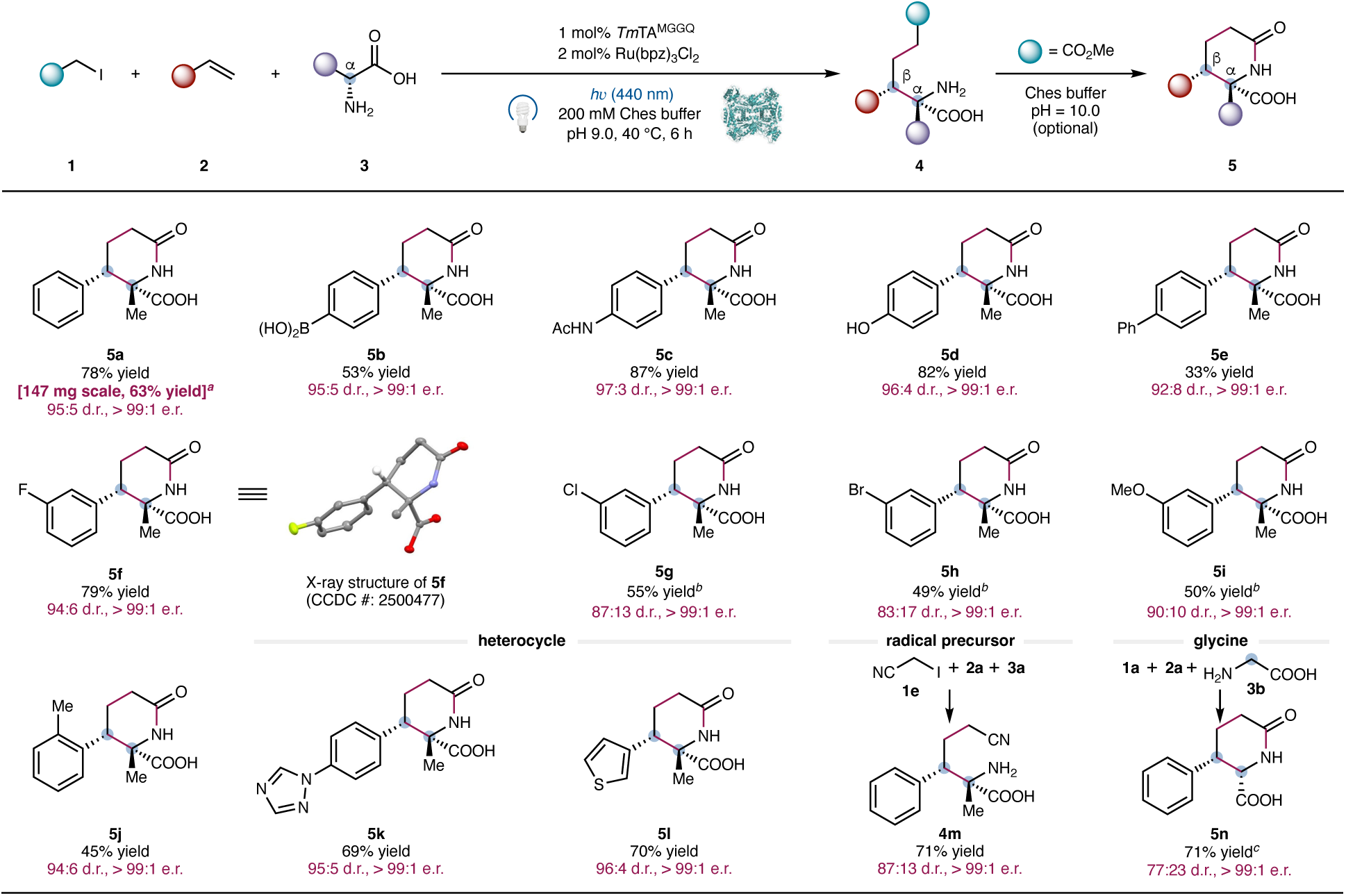
Substrate scope of the photobiocatalytic three-component coupling for the construction of contiguous tri- and tetrasubstituted stereocenters. Reaction conditions: **1** (2.0 equiv, 8.0 mM), **2** (1.0 equiv, 4.0 mM), **3** (5.0 equiv, 20.0 mM), 2 mol% Ru(bpz)_3_Cl_2_ (80 μM), 1 mol% *Tm*TA I84M W86G V29G K224Q (*Tm*TA^MGGQ^, 40 μM), 200 mM Ches buffer (pH = 9.0), *h*ν (440 nm Kessil LED, 40 W, 100% intensity), 40 °C, 6 h. Lactamization conditions: 400 μL Ches buffer (200 mM, pH = 10.0) was added, 60 °C, 6 h (see Supplementary Information for details). *^a^*0.5 mol% *Tm*TA^MGGQ^ was used. *^b^Tm*TA I84M W86G V29G (*Tm*TA^MGG^) was used in lieu of *Tm*TA^MGGQ^. *^c^Tm*TA E88V I84A F85R (*Tm*TA^VAR^) was used in lieu of *Tm*TA^MGGQ^.

### Directed evolution of biocatalysts for the construction of challenging adjacent tetrasubstituted stereocenters

The efficient and stereocontrolled construction of sterically congested contiguous tetrasubstituted stereocenters has remained a formidable challenge in asymmetric synthesis^29–31^. To address this limitation, we questioned whether our photobiocatalytic three-component coupling could be extended to access densely substituted amino acid derivatives bearing adjacent tetrasubstituted stereocenters. Using α-iodoacetate (**1a**), α-methylstyrene (**2o**), and _D_-alanine (_D_-**3a**) as the model substrates, we re-evaluated *Tm*TA variants from our collection in the presence of the optimal photocatalyst Ru(bpz)_3_Cl_2_ (Fig. 4a). When wild-type *Tm*TA was used, the desired three-component coupling product **5o** formed in only 2% yield, albeit with good diastereo- and enantioselectivity (92:8 d.r., >99:1 e.r., Fig. 4a, entry 1). Among the *Tm*TA variants examined (entries 2–6), *Tm*TA E88T exhibited the highest activity and diastereoselectivity, affording **5o** in 12% yield, 93:7 d.r., >99:1 e.r. (Fig. 4a, entry 4). Interestingly, the *Tm*TA I84M W86P variant, optimal for assembling contiguous tri- and tetrasubstituted stereocenters, showed substantially reduced activity in this context, indicating distinct active-site requirements for the formation of vicinal tetrasubstituted stereocenters *versus* mixed tri- and tetrasubstituted stereocenters. It was found that further increasing the photocatalyst loading to 2 mol% did not improve the yield of **5o**, indicating subtle reactivity differences between α-methylstyrene and styrene substrates (see Table S9 for details).

**Fig. 4.**
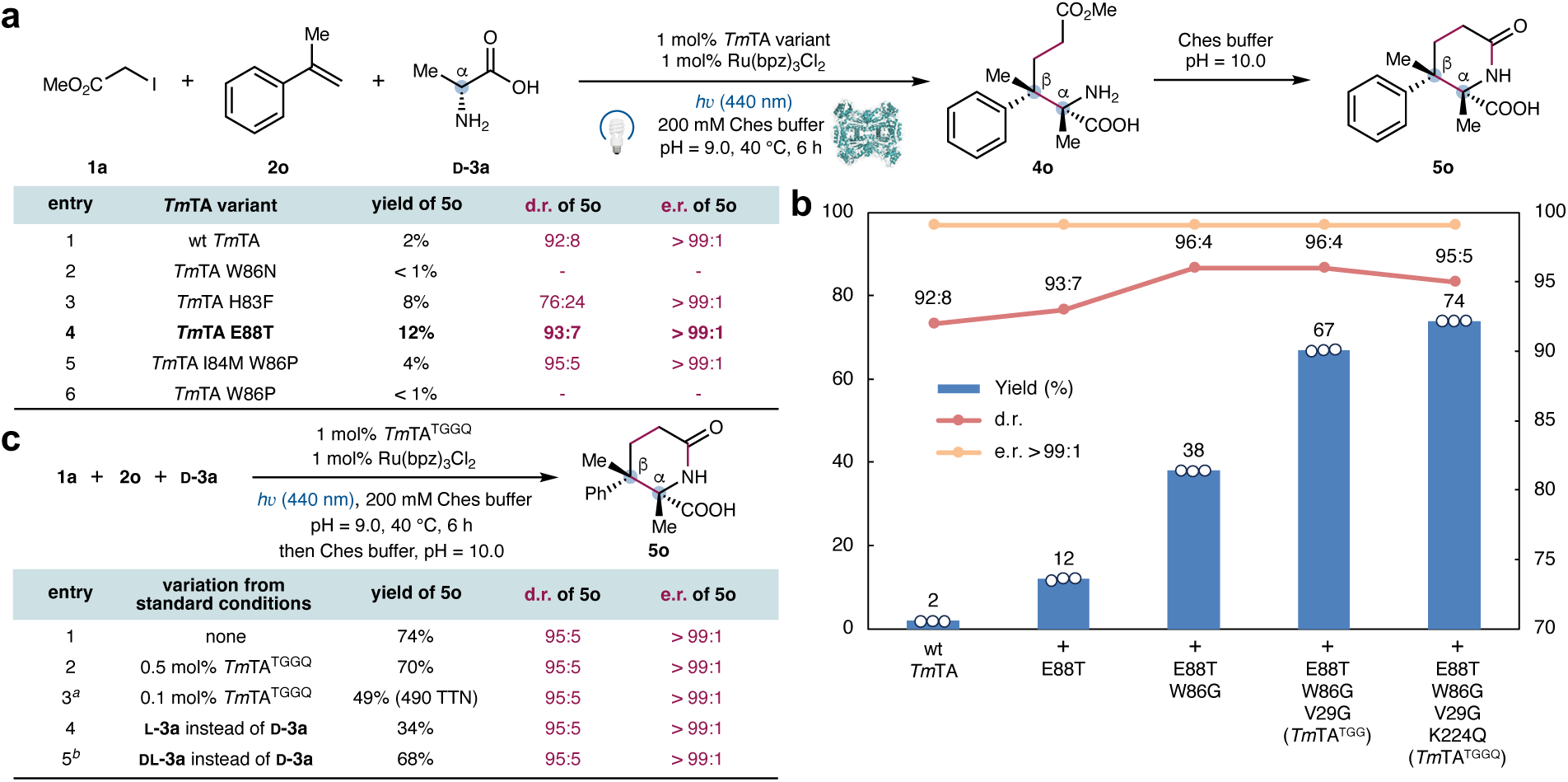
Evaluation and directed evolution of three-component coupling biocatalysts for the diastereo and enantioselective construction of challenging adjacent tetrasubstituted stereocenters. **a**, Evaluation of engineered *Tm*TA variants from in-house enzyme collection. Reaction conditions: **1a** (2.0 equiv, 32 mM), **2o** (1.0 equiv, 16 mM), **3a** (5.0 equiv, 80 mM), 1 mol% Ru(bpz)_3_Cl_2_ (160 μM), 1 mol% *Tm*TA variants (160 μM), 200 mM Ches buffer (pH = 9.0), *h*ν (440 nm, Kessil LED 40 W, 25% intensity), 40 °C, 6 h. Lactamization conditions: 400 μL Ches buffer (200 mM, pH = 10.0) was added, 60 °C, 6 h. **b**, Directed evolution of *Tm*TA E88T leading to the final variant *Tm*TA E88T W86G V29G K224Q (*Tm*TA^TGGQ^). **c**, Further evaluation of reaction conditions for photobiocatalytic three-component coupling. *^a^*0.3 mol% Ru(bpz)_3_Cl_2_ was used. *^b^*10 equiv _DL_-**3a** was used. TTN = total turnover number.

Encouraged by these results, we initiated a parallel directed evolution campaign using *Tm*TA E88T as the template to engineer highly active and stereoselective biocatalysts for construction contiguous tetrasubstituted stereocenters (Fig. 4b). Using the same photobiocatalytic screening platform, site-saturation mutagenesis and screening by targeting residue W86 identified a W86G substitution as a key beneficial mutation. The resulting *Tm*TA E88T W86G variant delivered a three-fold increase in activity, together with improved diastereoselectivity (38% yield, 96:4 d.r., >99:1 e.r.). Subsequent rounds of directed evolution focusing on residues V29 and K224 afforded a new quadruple mutant *Tm*TA E88T W86G V29G K224Q (*Tm*TA^TGGQ^), giving rise to **5o** in 74% yield, 95:5 d.r., and >99:1 e.r., corresponding to a 37-fold increase in activity relative to wild-type *Tm*TA. Further decreasing the enzyme loading to 0.5 mol% still afforded **5o** in 70% yield, 95:5 d.r. and >99:1 e.r. (Fig. 4c, entry 2). At 0.1 mol% enzyme loading, **5o** still formed in 49% yield, 95:5 d.r., >99:1 e.r. (Fig. 4c, entry 3), corresponding to a total turnover number (TTN) of 490. Comparison of the optimized variants evolved in the two contexts — *Tm*TA E88T W86G V29G K224Q for the construction of contiguous tetrasubstituted stereocenters and *Tm*TA I84M W86G V29G K224Q for the construction of contiguous tri- and tetrasubstituted stereocenters — revealed a striking degree of convergent evolution. Three key mutations, including W86G, V29G, and K224Q, were fully conserved across both evolution trajectories with different starting templates, suggesting that the critical role of these residues in accommodating the C–C bond forming radical addition step. Finally, both _L_- and _DL_-alanine (**3a**) were accepted by *Tm*TA^TGGQ^, leading to the same major stereoisomeric product with identical diastereo- and enantioselectivity (Fig. 4c, entries 4 and 5). Similar to the first model reaction, two-component coupling product **4o’** derived from **1a** and **2o** was also not observed.

### Substrate scope of photobiocatalytic construction of contiguous tetrasubstituted stereocenters

We next assessed the substrate scope of our evolved biocatalyst *Tm*TA^TGGQ^ to access sterically congested amino acid derivatives possessing contiguous tetrasubstituted stereocenters (Fig. 5). It was found that α-methylstyrenes bearing a sensitive *para*-substituent, including a methyl ester (**5p**), an acetamide (**5q**), and a hydroxy (**5r**) group, were smoothly transformed into the three-component coupling products with uniformly excellent stereocontrol. Heterocyclic substrates such as a pyrazole (**5s**), a thiophene (**5t**), and a furan (**5u**) were also found to be excellent coupling partners. Importantly, a range of α-substituents of the α-substituted vinylarene substrate, including an ethyl (**5v**), a methoxy (**5w**), a difluoromethyl (**5x**), a sensitive monofluoromethyl (**5y**), and a fluorine (**5z**) group, were compatible, further underscoring the synthetic utility of this three-component photobiocatalytic coupling for the synthesis of stereochemically well-defined organofluorine compounds^42^. In addition, 1-methyleneindane (**5aa**) underwent efficient conversion to deliver a spirocyclic product featuring adjacent tetrasubstituted stereocenters. Beyond olefinic coupling partners, *Tm*TA^TGGQ^ also accommodated a diverse array of radical precursors. The use of racemic α-methyl α-iodoacetate (**5ab**) afforded the product in 75% yield and >99:1 e.r., albeit with a nearly 1:1 diastereomeric ratio. α-Iodoketones (**6ac**) also represented excellent substrates, furnishing the corresponding cyclic imino acids in good yield with excellent diastereo- and enantiocontrol. Additional radical precursors, including α-iodoamides (**4ad**) and α-iodoacetonitriles (**4ae**), were also efficiently transformed to acyclic non-canonical amino acids featuring contiguous tetrasubstituted stereocenters with excellent yields and stereoselectivities. Finally, glycine (**5af**) was also readily accepted by *Tm*TA^VAR^, allowing the construction of adjacent tetra- and trisubstituted stereocenters. Thus, this photobiocatalytic platform enabled the collective stereoselective assembly of four types contiguous stereogenic centers (Fig. 1). The relative and absolute stereochemistry of products **5o** and **5z** were unambiguously established by single-crystal X-ray diffraction analysis (CCDC accession numbers 2500478 and 2500476, respectively, see the SI for further details).

**Fig. 5.**
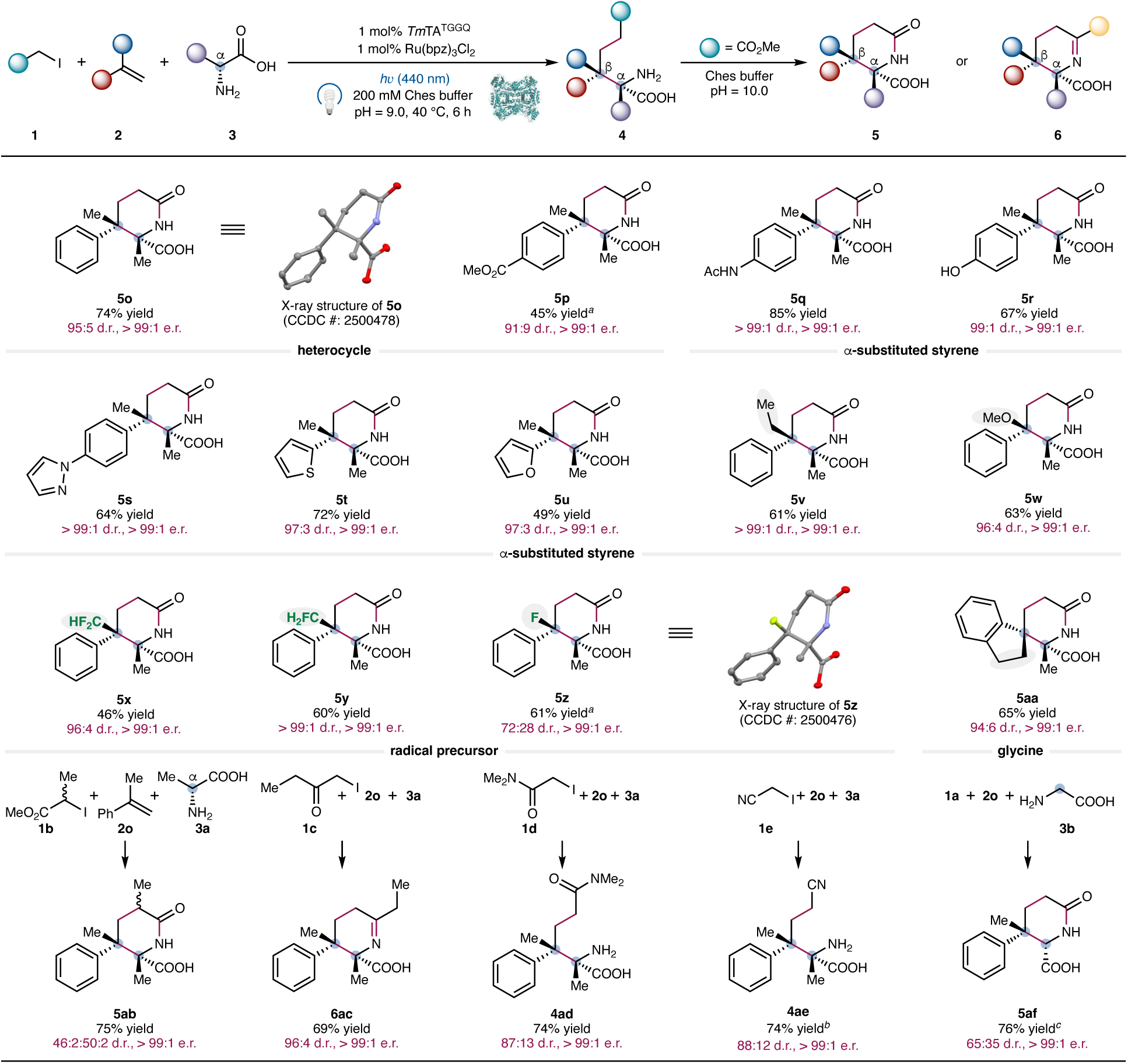
Substrate scope of the biocatalytic construction of two contiguous tetrasubstituted stereocenters. Reaction conditions were **1** (2.0 equiv, 32 mM), **2** (1.0 equiv, 16 mM), **3** (5.0 equiv, 80 mM), 1 mol% Ru(bpz)_3_Cl_2_ (160 μM), 1 mol% *Tm*TA E88T W86G V29G K224Q (*Tm*TA^TGGQ^, 160 μM), 200 mM Ches buffer (pH = 9.0), *h*ν (440 nm, Kessil LED 40 W, 25% intensity), 40 °C, 6 h. Lactamization conditions: 400 μL Ches buffer (200 mM, pH = 10.0) was added, 60 °C, 6 h (see Supplementary Information for details). *^a^Tm*TA E88T W86G V29G (*Tm*TA^TGG^) was used in lieu of *Tm*TA^TGGQ^. *^b^Tm*TA E88T W86G V29G K224C (*Tm*TA^TGGC^) was used in lieu of *Tm*TA^TGGQ^. *_c_Tm*TA E88V I84A F85R (*Tm*TA^VAR^) was used in lieu of *Tm*TA^TGGQ^.

### Preparative scale photobiocatalytic synthesis and downstream derivatization

To further demonstrate the synthetic utility of the current photobiocatalytic three-component coupling, we carried out preparative-scale reactions on a 1.0 mmol scale (Fig. 6). Photobiocatalytic coupling of **1a**, **2o** and **3a** on a 1.0 mmol scale furnished **5o** in 159 mg (64%) yield and excellent stereoselectivity. Sequential hydrolytic cleavage of **5o**, first under acidic conditions (3 M HCl) followed by basic treatment (6 M KOH), afforded the corresponding non-canonical amino acid **7** in 81% yield, 95:5 d.r., and >99:1 e.r. (Fig. 6a). Starting from **5o**, borane-mediated reduction provided the corresponding piperidine **8** bearing an α-hydroxymethyl group in 86% yield, 97:3 d.r., and >99:1 e.r. Given the prevalence of piperidines in pharmaceuticals^43^, this transformation highlights the ability of the present methods to provide streamlined access to polysubstituted piperidines featuring contiguous tetrasubstituted stereocenters. In addition, a second preparative photobiocatalytic reaction of **1c**, **2o** and **3a** on a 1.0 mmol (150 mg) scale delivered cyclic imino acid **6ac** in 58% yield, 96:4 d.r., and >99:1 e.r. (Fig. 6b). Subsequent reduction of **6ac** with NaBH_3_CN afforded piperidine **9**, which contained three stereocenters proximal to the N atom, in excellent yield and diastereoselectivity (92% yield, 95:5 d.r., >99:1 e.r.). The relative stereochemistry of piperidine **9** was established by 2D NOESY NMR spectroscopy (see the SI for details), which further supported the stereochemical assignment of enzymatic product **6ac**. Collectively, these preparative-scale reactions and downstream derivatization underscored the synthetic versatility and scalability of the stereochemically enriched products prepared through this photobiocatalytic three-component coupling platform.

**Fig. 6.**
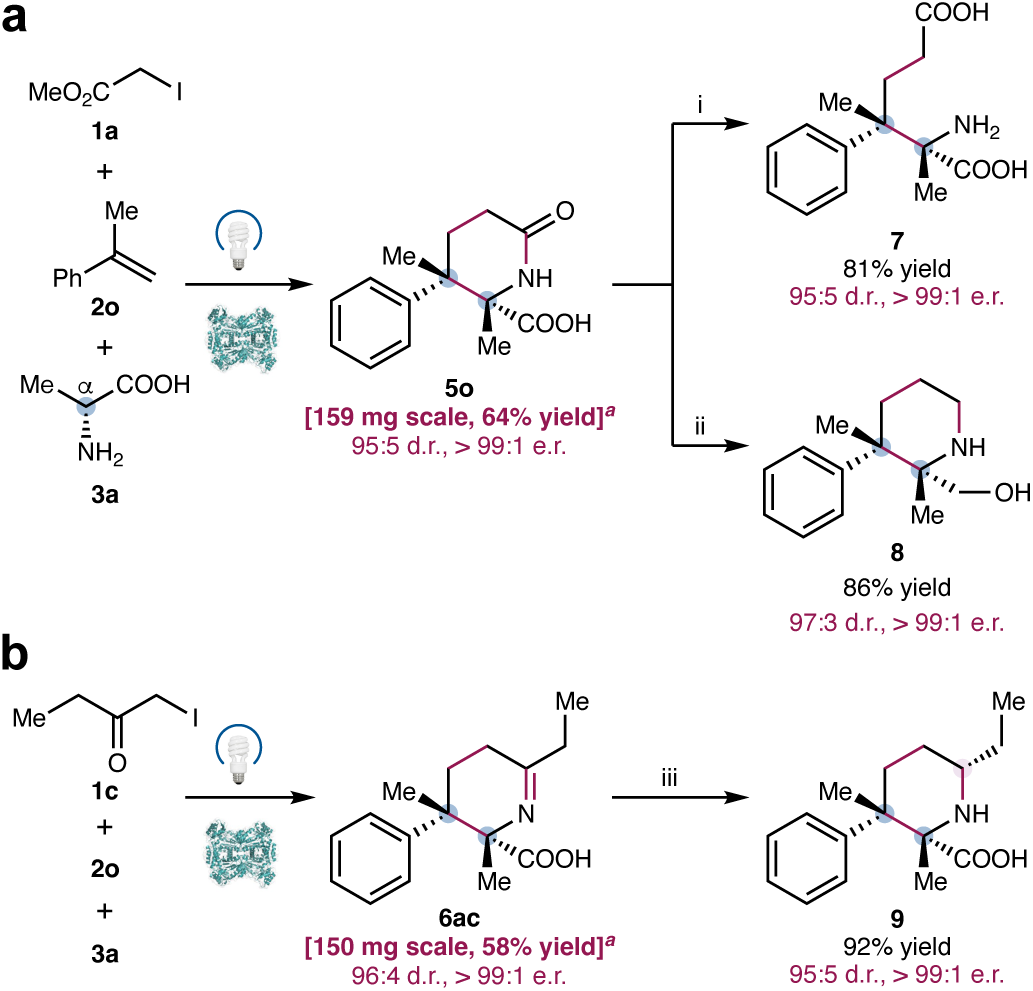
Preparative-scale photobiocatalytic synthesis and downstream chemical transformations. **a**, Preparative-scale photobiocatalytic synthesis of **5o** and derivatization to access stereochemically well-defined non-canonical amino acids and piperidines bearing two contiguous tetrasubstituted stereocenters. **b**, Preparative-scale photobiocatalytic synthesis of **6ac**, with subsequent transformations to obtain stereochemically well-defined piperidines possessing three stereocenters. Reaction conditions: i, 3 M HCl (two cycles), 70 °C, 12 h; 6 M KOH, 110 °C, 12 h; ii, BH_3_-THF, 75 °C, 24 h; iii, NaBH_3_CN, MeOH, 0 °C to rt, 6 h. *^a^*0.5 mol% *Tm*TA^TGGQ^ was used. See the Supplementary Information for details.

### Combinatorial photobiocatalytic synthesis in a library format

Due to the high synthetic versatility and broad substrate scope of our evolved biocatalysts, we applied the photobiocatalytic three-component coupling platform to combinatorial synthesis by simultaneously varying all three coupling partners. Five types of organic iodide radical precursors, including α-iodoacetate (**1a**), α-branched iodoacetate (**1b**), α-iodoketone (**1c**), α-iodoamide (**1d**) and α-iodonitrile (**1e**), for evaluation in conjunction with five distinct vinylarenes (**2a**, **2o**, **2x**, **2ag** and **2ah**) and two amino acid donors (alanine **3a** and glycine **3b**). The combinatorial biocatalytic synthesis was carried out in a single vessel in a 5 × 5 × 2 library format and each individual product was analyzed by UPLC-MS analysis. Of the 50 possible products, 40 had not been previously investigated in our substrate scope study (Figs. 3 and 5), representing 80% of the total library (structures of all 50 products are shown in Fig. S5). Chemoinformatic analysis revealed a high degree of structural diversity. Principal moments of inertia^44^ (PMI, I_1_, I_2_, and I_3_), calculated from force field-optimized structures using RDKit, indicated that the library spans a broad range of molecular shapes, with many compounds occupying regions sparsely populated among FDA-approved small-molecule drugs^45^, highlighting the structural novelty of the combinatorial library.

Using *Tm*TA^TGGQ^ as the biocatalyst, 96% of substrate combinations underwent successful photobiocatalytic coupling, providing reaction hits suitable for further optimization for medicinal chemistry applications^46^. Of these, 48% of products formed in >30% yield and 36% in 10–30% yield as determined with UPLC-MS analysis (Fig. 7a and Table S13). Glycine (**3b**) generally provided higher yields than alanine (**3a**) across this library with *Tm*TA^TGGQ^. In contrast, *Tm*TA^VAR^ (*Tm*TA E88V I84A F85R) showed complementary activity, providing higher overall yields with alanine (**3a**) (Fig. 7b and Table S13). Combining the two variants as a cocktail (0.5 mol% *Tm*TA^TGGQ^ + 0.5 mol% *Tm*TA^VAR^) further improved overall activity relative to single-component biocatalysts (Fig. 7c and Table S13). Using this strategy, 98% of the 5 × 5 × 2 library was synthesized, with 60% of compounds obtained in > 30% yield (Fig. 7d and Table S13). These results demonstrated the excellent potential of these three-component C–C coupling enzymes for combinatorial library synthesis.

**Fig. 7.**
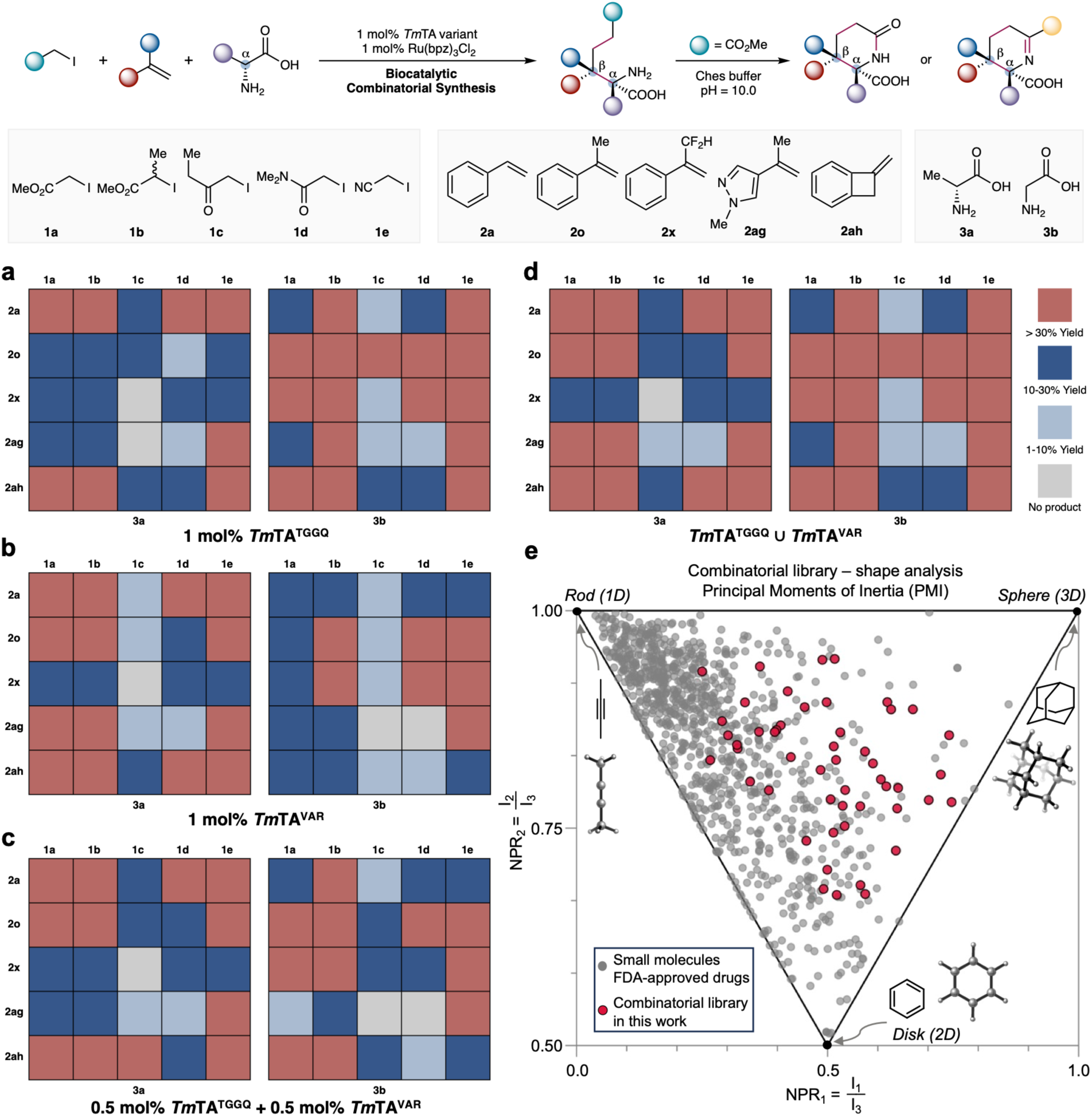
Photobiocatalytic combinatorial synthesis in a 5 × 5 × 2 library format. **a**, Heatmap of 5 × 5 × 2 photobiocatalytic combinatorial synthesis using 1.0 mol% *Tm*TA^TGGQ^. **b**, Heatmap of 5 × 5 × 2 photobiocatalytic combinatorial synthesis using 1.0 mol% *Tm*TA^VAR^. **c**, Heatmap of 5 × 5 × 2 photobiocatalytic combinatorial synthesis using a cocktail of two variants (0.5 mol% *Tm*TA^TGGQ^ + 0.5 mol% *Tm*TA^VAR^). **d**, Coverage of 98% products in the 5 × 5 × 2 library using *Tm*TA^TGGQ^ and *Tm*TA^VAR^ (yield shown corresponded to the best-performing variant). **e**, Normalized principal moment of inertia (PMI) analysis of molecular shapes in the library. NPR: Normalized PMI ratios. Reaction conditions: **1** (2.0 equiv, 32 mM), **2** (1.0 equiv, 16 mM), **3** (5.0 equiv, 80 mM), 1 mol% Ru(bpz)_3_Cl_2_ (160 μM), 1 mol% *Tm*TA variants (160 μM), 200 mM Ches buffer (pH = 9.0), *h*ν (440 nm, Kessil LED 40 W, 25% intensity), 40 °C, 6 h. Lactamization conditions: 400 μL Ches buffer (200 mM, pH = 10.0) was added, 60 °C, 6 h (see Supplementary Information for details).

### Mechanistic and computational studies

To gain insights into the mechanism of these photobiocatalytic three-component coupling processes, we performed radical trapping experiments using 2,2,6,6-tetramethyl-1-piperidinyloxy (TEMPO, Fig. 8a). For the reaction of **1a**, **2a** and **3a**, the addition of 4 equiv TEMPO completely suppressed formation of the desired C–C coupling product **5a**. Instead, two radical trapping products were observed: **10**, derived from the α-carbonyl radical generated from **1a**, and **11**, derived from the secondary benzylic radical formed by radical addition to styrene **2a**, in 41% and 13% yield, respectively, as determined by ^1^H NMR spectroscopic analysis using an internal standard. When α-methylstyrene **2o** was used in place of **2a**, a slightly lower yield of 12% was observed for **10**. TEMPO-trapping product **12** generated from the tertiary benzylic radical was also observed by LC-MS analysis, although its yield could not be accurately determined due to limited stability. To our knowledge, this represented the first observation of both TEMPO-trapped radical intermediates in a biocatalytic three-component coupling, providing direct evidence for the proposed radical mechanism of the transformation.

**Fig. 8.**
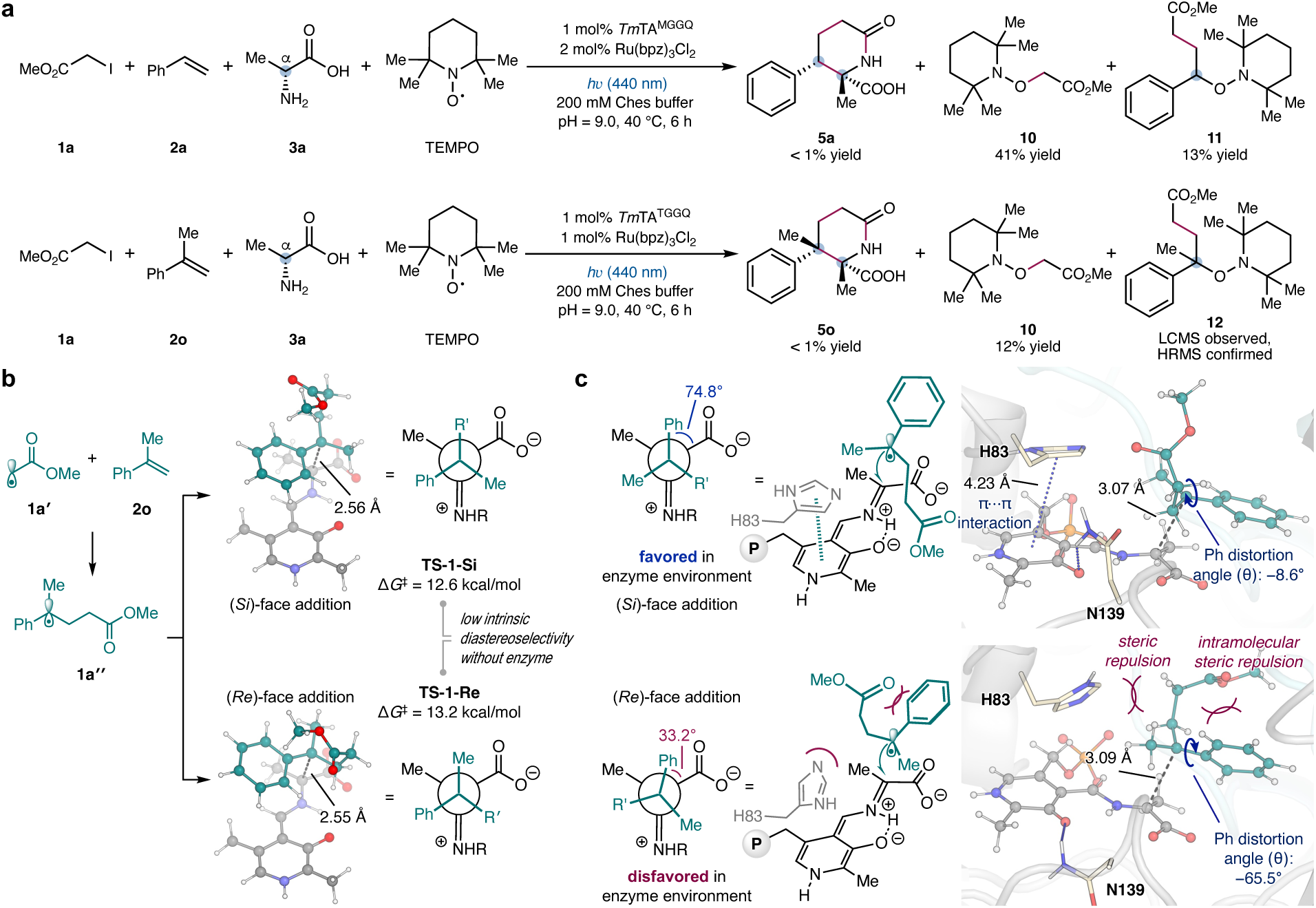
Mechanistic and computational studies on the enzymatic three-component radical coupling. **a**, TEMPO trapping studies. **i**, Reaction conditions: **1a** (2.0 equiv, 8.0 mM), **2a** (1.0 equiv, 4.0 mM), **3a** (5.0 equiv, 20.0 mM), 400 mol% TEMPO (16.0 mM), 1 mol% *Tm*TA^MGGQ^ (40 μM), 2 mol% Ru(bpz)_3_Cl_2_ (80 μM), 200 mM Ches buffer (pH = 9.0), *h*ν (440 nm Kessil LED, 40 W, 100% intensity), 40 °C, 6 h. **ii**, Reaction conditions: **1a** (2.0 equiv, 32 mM), **2o** (1.0 equiv, 16 mM), **3a** (5.0 equiv, 80 mM), 400 mol% TEMPO (64 mM), 1 mol% *Tm*TA^TGGQ^ (160 μM), 1 mol% Ru(bpz)_3_Cl_2_ (160 μM), 200 mM Ches buffer (pH = 9.0), *h*ν (440 nm, Kessil LED 40 W, 25% intensity), 40 °C, 6 h. **b**, DFT-optimized structures of radical addition transition states with α,α-disubstituted benzyl radical **1a’’** (**TS-1-*Si*** and **TS-1-*Re***) using a truncated PLP cofactor model showing low intrinsic diastereoselectivity in radical addition in the absence of active site residues. Gibbs free energies of activation (Δ*G*^‡^) are relative to the quinonoid species and the radical intermediate. **c**, Near-attack conformations (NACs) for the radical addition pathways from classical MD simulations. The Newman projections illustrate the conformations about the forming C–C bond in the radical addition, highlighting the effect of the enzyme environment observed in NAC simulations.

To further investigate the origin of the high diastereoselectivity in the enzymatic three-component radical coupling, we performed computational studies using density functional theory (DFT) calculations and classical molecular dynamics (MD) simulations. Using a truncated PLP model, DFT calculations revealed that the reaction between the quinonoid intermediate and the (*Si*)-face of the prochiral α,α-disubstituted benzyl radical is favored over the (*Re*)-face of the radical by 0.6 kcal/mol (Fig. 8b). The low intrinsic diastereoselectivity with this PLP model arises from the lack of steric contributions provided by the active site residues. Classical MD simulations were then carried out to model near-attached conformations (NACs) for the addition to the (*Si*)- and (*Re*)-faces of the α,α-disubstituted benzyl radical, by restraining the distance between the benzylic radical carbon and the quinonoid Cα atom to 2.7–3.1 Å, based on the DFT-computed transition-state geometries. In the (*Si*)-face addition NAC (Fig. 8c), the benzylic radical and the quinonoid adopt a staggered conformation about the forming C–C bond, minimizing unfavorable steric clashes. In contrast, in the (*Re*)-face addition NAC, steric repulsion between residue H83 and its alkyl substituent (R′) distorts the benzylic radical, producing a large dihedral angle about the C(Ph)–Cα bond (−65.5°) and preventing π–π interactions between H83 and the quinonoid intermediate. These unfavorable steric interactions collectively disfavor this (*Re*)-face addition pathway.

To gain further insights into the beneficial mutation effects, classical MD simulations were performed to model the NAC for C–C bond formation from the (*Si*)-face of the α,α-disubstituted benzyl radical in both wt *Tm*TA and evolved *Tm*TA^TGGQ^. In wt *Tm*TA, a cation/π interaction between the guanidium group of R140 and the indole side chain of W86 blocks the approach of the benzyl radical to the PLP covalent intermediate, leading to increased steric repulsion in the NAC (Extended Data Fig. 1). This effect is reflected by the larger distortion angle, θ, of the phenyl group on the benzyl radical This steric repulsion along the substrate tunnel is relieved in *Tm*TA^TGGQ^ by the beneficial W86G mutation, as indicated by a smaller phenyl distortion angle and a more favorable radical geometry in the NAC for the C–C bond formation.. Furthermore, in wt *Tm*TA, an internal hydrogen bonding between the carboxylate of E88’s and K224’s ε-NH_3+_ impedes the radical access to the PLP intermediate. By introducing E88T and K224Q mutations into wt *Tm*TA, the E88–K224 hydrogen bond is disrupted in *Tm*TA^TGGQ^, thereby increasing the flexibility of the 82–86 loop and further expanding the active-site volume. The expanded active site is evidenced by an increased solvent-accessible surface area of the PLP intermediate from 35.8 Å^2^ in wt *Tm*TA to 46.0 Å^2^ in *Tm*TA^TGGQ^. Furthermore, as a key beneficial mutation, V29G substantially expands the substrate access tunnel (Figure S23) by eliminating the bulky side chain at residue 29. Collectively, all these beneficial mutations in *Tm*TA^TGGQ^ expand the substrate entrance tunnel, allowing more efficient transfer of the benzyl radical and facilitating radical C–C coupling within the enzyme active site.

**Extended Data Fig. 1.**
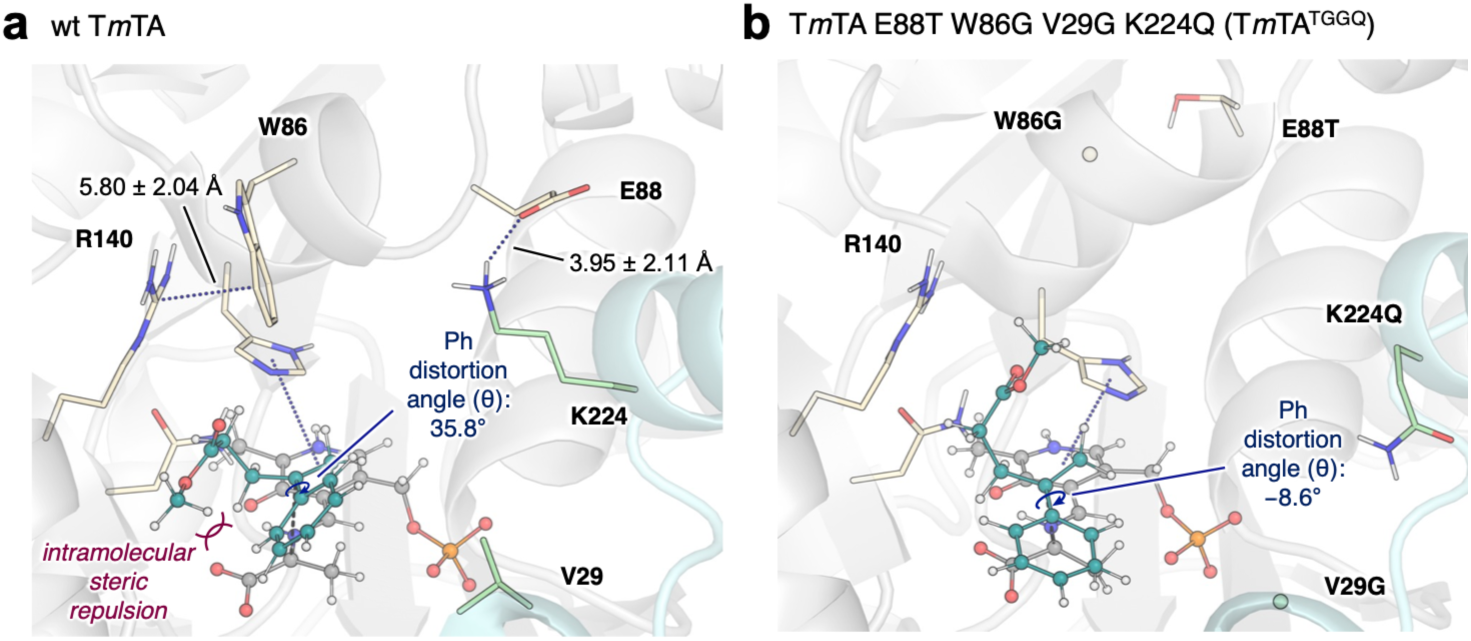
Effects of beneficial mutations introduced to *Tm*TA^TGGQ^. **a,** Near-attack conformations (NACs) from classical MD simulations for the radical-mediated C–C bond formation in wt *Tm*TA. **b,** Near-attack conformations (NACs) from classical MD simulations for the radical-mediated C–C bond formation in *Tm*TA^TGGQ^.

## Conclusion

In summary, we developed a novel photobiocatalytic three-component C–C coupling via a C–H functionalization strategy. Directed evolution of thermophilic threonine aldolases allowed the diastereo- and enantioselective construction of highly challenging contiguous stereocenters, including adjacent tri- and tetrasubstituted stereocenters and two contiguous tetrasubstituted stereocenters, addressing a long-standing problem in asymmetric radical catalysis. This approach provided access to a range of valuable stereochemically complex non-canonical amino acid derivatives and nitrogen heterocycles ubiquitous in medicinally important agents. The broad substrate scope of our evolved biocatalysts further enabled the diversity-oriented synthesis of valuable products in a combinatorial library format. Collectively, these stereoselective three-component couplings represented some of the most complex non-native biocatalytic reactions reported to date and constitute a rare example of enzymatic trimolecular reactions previously unknown in both synthetic organic chemistry and native enzymology. We anticipate that this study will inspire the development of novel biocatalytic multicomponent reactions to tackle challenging problems beyond the reach of conventional asymmetric catalysis.

## Methods

### Expression of engineered *Tm*TA variants

A single colony from LB_kan_ agar plate was picked using a sterile toothpick and cultured in LB_kan_ media (25 mL) at 37 °C and 230 rpm overnight. This preculture was used to inoculate 0.6 L of TB_kan_ media (v/v = 2.0%) in a 2 L Erlenmeyer flask, and the expression culture was incubated at 37 °C and 230 rpm for ca. 3 h until the OD_600_ reached ca. 2.0. The culture was cooled on ice for 30 min and induced with 0.5 mM isopropyl-*β*-_D_-thiogalatopyranoside (IPTG) (final concentration). Protein expression was conducted at 22 °C and 200 rpm for 20 h. *E. coli* cells were harvested by centrifugation at 4 °C and 6,000 *g* for 10 min using a Thermo Scientific Sorvall Lynx 6000 superspeed centrifuge. The expression levels of *Tm*TA^MGGQ^ and *Tm*TA^TGGQ^ were 90 mg from 0.6 L TB culture (150 mg/L), and 110 mg from 0.6 L TB culture (183 mg/L), respectively.

### Analytical scale photobiocatalytic three-component radical coupling using the construction of two contiguous tetrasubstituted stereocenters as an example

The threonine aldolase variant *Tm*TA^TGGQ^ was allowed to thaw and kept on top of ice. In a Coy anaerobic chamber, the following stock solutions were prepared: α-substituted styrene substrate (200 mM in degassed DMSO), radical precursor substrate (400 mM in degassed DMSO), amino acid substrate (1000 mM in degassed 200 mM Ches buffer, pH = 9.0) and Ru(bpz)_3_Cl_2_ (2.0 mM in degassed DMSO). To a Fisher brand reaction tube (13 **×** 100 mm, 8 mL) containing a stir bar were added Ches buffer (200 μL, 200 mM, pH = 9.0), 30 μL α-substituted styrene substrate stock solution, 30 μL radical precursor substrate stock solution, 30 μL amino acid substrate stock solution, 30 μL Ru(bpz)_3_Cl_2_ stock solution and ca. 40 μL *Tm*TA^TGGQ^ solution (the volume of the enzyme stock solution is dependent on the concentration of the specific enzyme sample). The reaction tube was then sealed and removed from the Coy anaerobic chamber and submerged in a water bath at 40 °C, where it was allowed to stir at 250 rpm and 40 °C and illuminated with two 40 W Kessil LED lamps (440 nm, 25% intensity) for 6 h. For lactam formation, Ches buffer (400 μL, 200 mM, pH = 10.0) was added and the mixture was heated at 60 °C for another 6 h. The biocatalytic reaction was then worked up and analyzed by HPLC-MS analysis.

### Preparative scale photobiocatalytic three-component radical coupling using the construction of two contiguous tetrasubstituted stereocenters as an example

The threonine aldolase variant *Tm*TA^TGGQ^ was allowed to thaw and kept on top of ice. In a Coy anaerobic chamber, the following stock solutions were prepared: α-substituted styrene (200 mM in degassed DMSO), radical precursor (400 mM in degassed DMSO), amino acid substrate (1000 mM in degassed 200 mM Ches buffer, pH = 9.0) and Ru(bpz)_3_Cl_2_ (2.0 mM in degassed DMSO). To a 50 mL round-bottom flask containing a stir bar were added Ches buffer (7 mL, 200 mM, pH = 9.0), 1.0 mL α-substituted styrene stock solution, 1.0 mL radical precursor stock solution, 1.0 mL amino acid substrate stock solution, 1.0 mL Ru(bpz)_3_Cl_2_ stock solution and ca. 1.0 mL *Tm*TA^TGGQ^ stock solution. The round-bottom flask was then sealed with a rubber septum and vinyl tapes and removed from the Coy anaerobic chamber. The reaction mixture was allowed to stir at 450 rpm and 40 °C in a water bath and illuminated with two 40 W Kessil LED lamps (440 nm, 25% power output, distance of Kessil LED to reaction flask = 8 cm) for 12 h. For lactam formation, 15 mL Ches buffer (200 mM, pH = 10.0) was added and the mixture was heated at 60 °C for another 6 h. The biocatalytic reaction was then worked up and purified by flash column chromatography with the aid of a Biotage Selekt.

## Acknowledgements

This research is supported by the National Institutes of Health (R01EB036084). Transformations of nitrogen heterocycles is supported by the Army Research Office (W911NF-24-2-0246). The computational study is supported by the National Science Foundation (CHE-2400087). Additional support on enzyme mining is provided by the Herman Frasch Foundation (947-HF22). Y.Y. is an Alfred P. Sloan Research Fellow (FG-2024-22244), a Camille Dreyfus Teacher-Scholar Awardee (TC-25-084), a David & Lucile Packard Fellow (2023-76169) and a Howard Hughes Medical Institute Freeman Hrabowski Scholar. Molecular dynamics simulations were performed at the Center for Research Computing of the University of Pittsburgh and the Advanced Cyberinfrastructure Coordination Ecosystem: Services & Support (ACCESS) program supported by the National Science Foundation grant numbers OAC-2117681 and OAC-2138259. We thank Dr. Tian-Ci Wang (University of California Santa Barbara) for helpful discussions and Dr. Quanquan Wang (University of California Santa Barbara) for preparing **1b**, **1c** and **1d**.

## Data availability

All data are available in the main text and the Supplementary Information. Plasmids encoding evolved *Tm*TA variants reported in this study are available for research purposes from Y.Y. under a material transfer agreement with the University of California Santa Barbara and Howard Hughes Medical Institute.

## Author contributions

Y.Y. conceived and directed the project. Q.H. and C-X.Z. designed and performed the experimental studies. B.K.M. carried out the computational studies. Q.H., B.K.M., P.L. and Y.Y. wrote the manuscript with the input of all other authors.

## Competing interests

Y.Y. and Q.H. are inventors on a patent application submitted by the University of California Santa Barbara and Howard Hughes Medical Institute that covers compositions, systems, and methods for photobiocatalytic three-component coupling using threonine aldolases. The remaining authors declare no competing interests.

## Notes

### Competing Interest Statement

The authors have declared no competing interest.

